# *TP53* Inactivation Confers Resistance to the Menin Inhibitor Revumenib in Acute Myeloid Leukemia

**DOI:** 10.1101/2025.03.24.644993

**Authors:** Jeevitha D’Souza, Camille J. Leung, Aishwarya C. Ballapuram, Abigail S. Lin, Alexa Rane Batingana, Adam J. Lamble, Rhonda E. Ries, Carolina E. Morales, Charisa Cottonham, Pranav Gona, Li Fan, Xiaotu Ma, Kevin Shannon, Soheil Meshinchi, Benjamin S. Braun, Benjamin J. Huang

**Affiliations:** Department of Pediatrics, University of California San Francisco, San Francisco, CA, USA; Division of Hematology and Oncology, Seattle Children’s Hospital, Seattle, WA, USA; Fred Hutchinson Cancer Center, Seattle, WA, USA; Department of Computational Biology, St. Jude Children’s Research Hospital, Memphis, TN, USA; Helen Diller Family Comprehensive Cancer Center, University of California San Francisco, San Francisco, CA, USA

## Abstract

Acute myeloid leukemia (AML) is a heterogeneous cancer that is associated with poor outcomes. Revumenib and other menin inhibitors have shown promising activity against AMLs with *KMT2A*-rearrangements or *NPM1* mutations. However, mechanisms of *de novo* resistance have not yet been elucidated. We analyzed a panel of cell lines and generated an isogenic model to assess the impact of *TP53* mutations on the response of AML cells to revumenib. *TP53* mutations are associated with *de novo* resistance to revumenib, impaired induction of *TP53* transcriptional targets, and deregulated expression of the BH3 proteins BCL-2 and MCL-1. The MCL-1 inhibitor MIK665, but not venetoclax, preferentially sensitized *TP53*-mutant AML cells to revumenib. These data identify mutant *TP53* as a potential biomarker for *de novo* resistance to revumenib, and provide a rationale to evaluate MCL-1 and menin inhibitor combinations in patients *KMT2A*-rearranged leukemias with *TP53* mutations.

## INTRODUCTION

Menin is an essential transcriptional cofactor for the initiation and maintenance of *KMT2A*-rearranged leukemias^1^. This landmark discovery stimulated drug discovery efforts^2–6^ that resulted in the development of revumenib^7^ and other menin inhibitors^8,9^. Revumenib showed promising activity in early phase clinical trials^10^, resulting in recent FDA approval for patients with relapsed, refractory *KMT2A*-rearranged leukemias. While on target *MEN1* mutations are a common mechanism of adaptive resistance to menin inhibitors^11^, mutations that confer *de novo* resistance have not been elucidated.

*TP53* mutations are associated with poor outcomes to standard chemotherapy regimens^12^ and to newer agents such as venetoclax^13–15^ in patients with acute myeloid leukemia (AML). While the hypomethylating agent decitabine showed activity in *TP53*-mutated AML, these responses were not durable^16^ and subsequent studies revealed less robust response rates^17,18^. Clinical trials of menin inhibitors reported to date have not accrued enough patients to address how *TP53* mutational status modulates treatment responses.

Here we show that *KMT2A*-rearranged AML cell lines that harbor *TP53* mutations exhibit *de novo* resistance to the menin inhibitor revumenib. These data were validated in an isogenic *TP53* cell line model. Furthermore, *TP53* mutational status modulated the transcriptional response of *KMT2A*-rearranged AML cells to revumenib at the level of *TP53* regulated gene sets, resulting in differential regulation of BCL-2 and MCL-1. Accordingly, the MCL-1 inhibitor MIK665 potently sensitizes *TP53*-mutant AML cells to revumenib. These data have implications for identifying AML patients who are at high risk of failing menin inhibitor treatment and for developing therapeutic strategies for overcoming resistance.

## RESULTS

### *KMT2A*-rearranged AML cell lines with *TP53* mutations exhibit *de novo* resistance to revumenib

We exposed *KMT2A*-rearranged (THP-1, NOMO-1, and MV-4-11, **Fig. S1**) and *NPM1*-mutated (OCI-AML3) AML cell lines to varying doses of revumenib. Experiments that utilized CellTiter-Glo (CTG) luminescence as an endpoint measure of viable cells revealed that both *TP53*-mutant cell lines (THP-1 and NOMO-1) were more resistant to revumenib than the two AML cell lines with wildtype (WT) *TP53* (OCI-AML3 and MV-4-11) (**Fig. 1A and 1B**). We performed similar experiments utilizing cleaved caspase-3 (CC3) detection as a readout for apoptosis. Remarkably, whereas revumenib resulted in a dose dependent increase in CC3 staining in *TP53* WT OCI-AML3 and MV-4-11 cells, THP-1 or NOMO-1 cells were completely resistant to apoptosis (**Fig. 1C**). By contrast, all four AML cell lines robustly induced CC3 expression in response to daunorubicin (**Fig. 1D**). Whereas exposing *TP53* WT MV-4-11 and OCI-AML3 cells to varying doses of revumenib for 96 hours resulted in a dose dependent increase in CD68 and CD11B cell surface antigen expression, *TP53*-mutant THP-1 and NOMO-1 cells failed to upregulate either myeloid differentiation marker (**Fig. 1E and 1F**).

**Figure 1.**
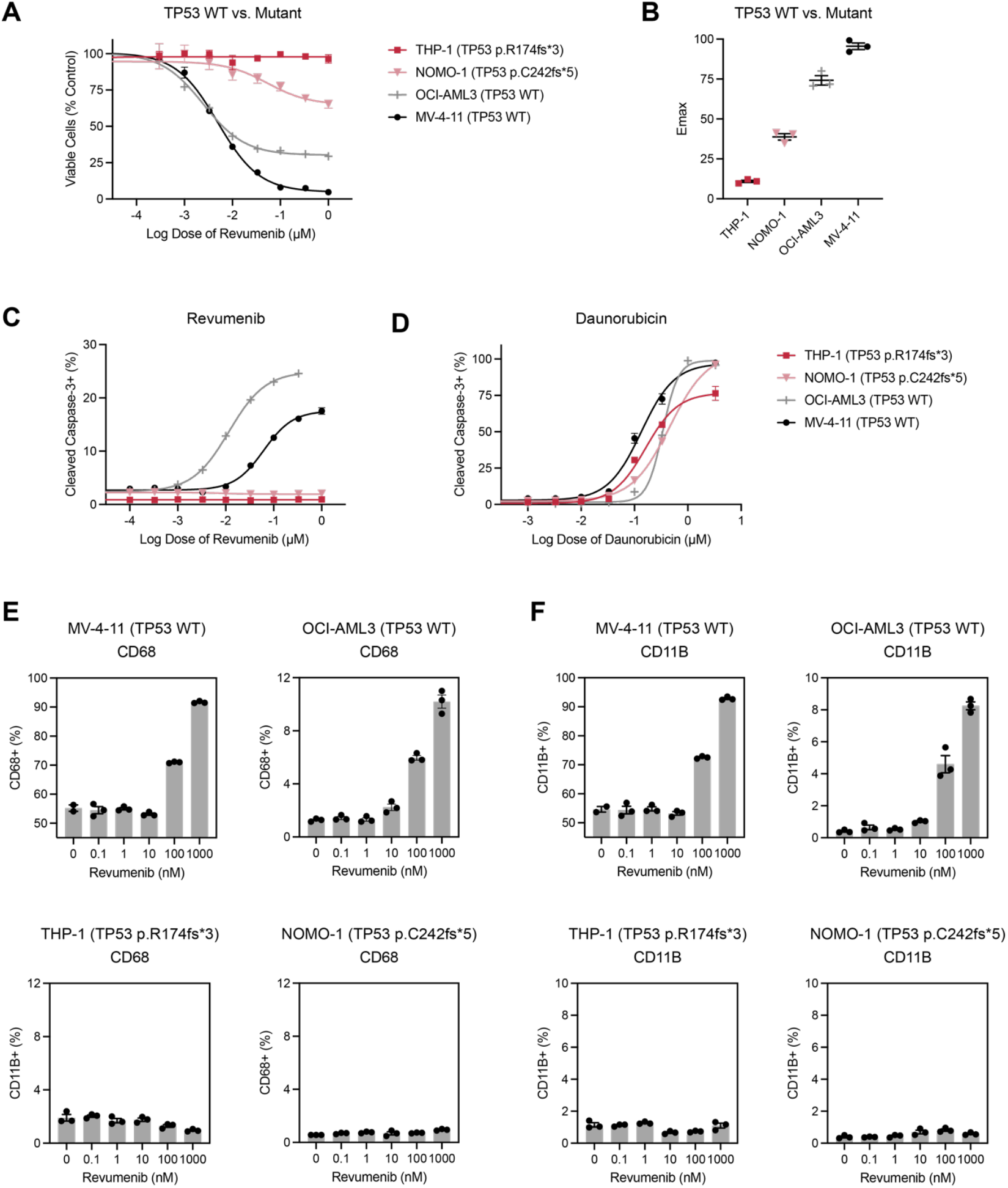
Activity of revumenib in *TP53* wildtype versus *TP53*-mutant AML cell lines. **A.** Dose response relationship of THP-1 (*TP53* p.R174fs*3), NOMO-1 (*TP53* p.C242fs*5), OCI-AML3 (*TP53* wildtype or WT), and MV-4-11 (*TP53* WT) cells after exposure to single agent revumenib for 72 hours. **B.** Maximal effect (E_max_) of revumenib in cell lines over 3 independent experiments each performed in technical triplicate. **C. and D.** Percentage of cells positive for cleaved caspase-3 after exposure to single agent **C.** revumenib or **D.** daunorubicin for 48 hours. **E.** CD68 and **F.** CD11B cell surface protein expression for AML cells after exposure to varying doses of revumenib for 96 hours.

### *TP53* inactivation causes revumenib resistance in AML cell lines

To directly ask if *TP53* inactivation confers resistance to revumenib in an isogenic cell line model, we performed CRISPR/Cas9 gene editing to generate MV-4-11 cells harboring a homozygous ~1.2 kilobase deletion in *TP53* (**Fig. S2**). Exposing *TP53* WT and *TP53* knockout (KO) MV-4-11 cell lines to varying doses of revumenib revealed that induction of CC3 was markedly impaired in *TP53* KO MV-4-11 cells versus *TP53* WT MV-4-11 cells (**Fig. 2A**). *TP53* inactivation also reduced the apoptotic response to daunorubicin in this model (**Fig. 2B**). Additionally, revumenib-induced myeloid differentiation was blunted in *TP53* KO MV-4-11 cells (**Fig. 2C**). Since clinical trial development is currently focused on testing revumenib in combination with standard chemotherapy, we performed Bliss independence synergy analysis on *TP53* WT and *TP53* KO MV-4-11 cells to assess apoptosis over a range of revumenib and daunorubicin doses. While this combination potentiated CC3 induction and exhibited positive synergy scores in *TP53* WT MV-4-11 cells (**Fig. 2D**), no synergy was observed in *TP53* KO MV-4-11 cells (**Fig. 2E**). Similar results were observed in other AML cell lines treated with the revumenib and daunorubicin combination (**Figs. 2F**).

**Figure 2.**
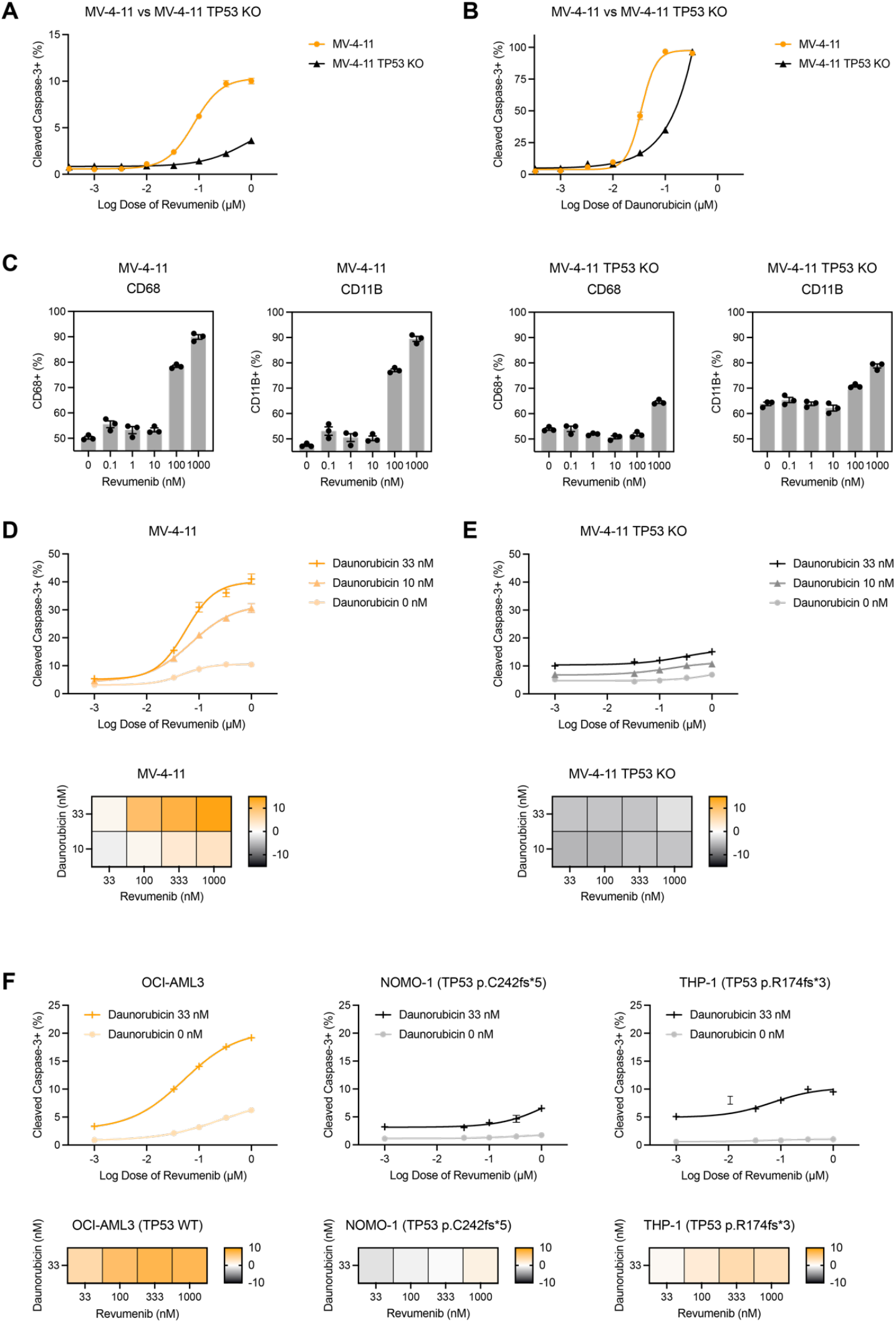
Synergy of revumenib and daunorubicin in isogenic *TP53* wildtype versus knockout AML cells. **A. and B.** Percentage of cells positive for cleaved caspase-3 (CC3+) with MV-4-11 *TP53* wildtype (WT) versus *TP53* knockout (KO) after exposure to single agent **A.** revumenib or **B.** daunorubicin for 48 hours. **C.** CD68 and CD11B cell surface protein expression for MV-4-11 *TP53* WT versus *TP53* KO cells after exposure to varying doses of revumenib for 96 hours. **D. and E.** CC3+ dose response of **D.** MV-4-11 *TP53* WT versus **E.** *TP53* KO after exposure to single agent revumenib versus revumenib combined with daunorubicin. Bliss synergy scores with various doses of revumenib and daunorubicin in MV-4-11 *TP53* WT versus *TP53* KO cells are shown in the panels below each dose response experimental result. **F.** CC3+ dose response for OCI-AML3 (*TP53* wildtype), NOMO-1 (*TP53* p.C242fs*5), or THP-1 (*TP53* p.R174fs*3) cells after exposure to single agent revumenib versus revumenib combined with daunorubicin for 48 hours, with associated Bliss synergy scores.

### *TP53* inactivation results in altered transcriptional response to revumenib

To characterize potential mechanisms of *de novo* to revumenib resistance in *TP53*-mutant AML, we exposed *TP53* WT and *TP53* KO MV-4-11 cells to either DMSO or 1000 nM of revumenib for 24 hours and performed bulk RNA sequencing (RNA-seq). Interestingly, gene set enrichment analysis (GSEA) focused on *HOXA9-MEIS1* gene sets showed that *TP53* WT and *TP53* KO cells both exhibited potent downregulation of *HOXA9-MEIS1* target genes in response to revumenib (**Fig. 3A**). By contrast, GSEA focused on *TP53* gene sets revealed that *TP53* WT MV-4-11 cells significantly upregulated *TP53* transcriptional pathways compared to *TP53* KO MV-4-11 cells (**Fig. 3B**). Additionally, BH3 family members such as MCL-1 and BCL-2 were differentially expressed (**Fig. S3**), implicating them as potential mediators of apoptosis resistance in *TP53*-mutant AML.

**Figure 3.**
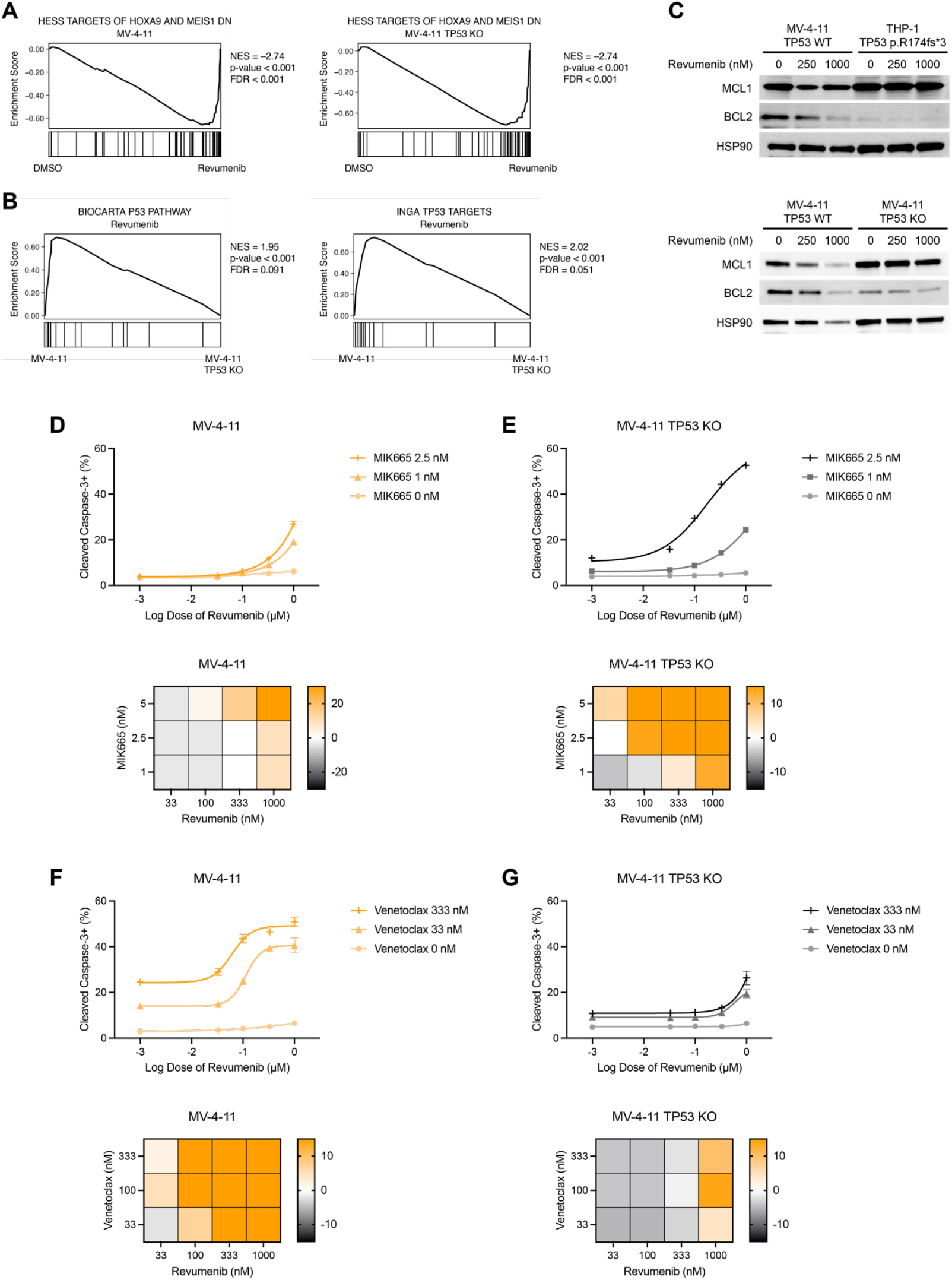
Impact of revumenib on transcriptional response and combining revumenib with BCL-2 and MCL-1 inhibitors. **A.** Gene set enrichment analysis (GSEA) of MV-4-11 *TP53* WT and MV-4-11 *TP53* KO cells, comparing treatment with DMSO versus 1000 nM revumenib for 24 hours, focused on genes down-regulated in hematopoietic precursor cells conditionally expressing *HOXA9* and *MEIS1*. **B.** GSEA comparing MV-4-11 *TP53* WT versus *TP53* KO cells treated with revumenib, focused on *TP53* pathway and target genes. **C.** Western blot of MCL-1 and BCL-2 in MV-4-11 versus THP-1 cells (top panel) and isogenic MV-4-11 *TP53* wildtype (WT) versus *TP53* knockout (KO) cells (bottom panel), treated with 0, 250, and 1000 nM of revumenib for 48 hours. **D. and E.** Percentage of cells positive for cleaved caspase-3 (CC3+) with **D.** MV-4-11 *TP53* WT versus **E.** *TP53* KO after exposure to single agent revumenib versus revumenib combined with the MCL-1 inhibitor MIK665. Bliss synergy scores with various doses of revumenib and venetoclax are shown in the panels below each dose response experimental result. **F. and G.** CC3+ dose response with **G.** MV-4-11 *TP53* WT versus **G.** *TP53* KO after exposure to single agent revumenib versus revumenib combined with venetoclax, with associated Bliss synergy scores.

### MCL-1 inhibition resensitizes *TP53* KO MV-4-11 cells to revumenib

Whereas Western blotting showed that *TP53* WT MV-4-11 cells downregulated MCL-1 and BCL-2 protein levels after revumenib exposure, treatment had no effect on MCL-1 expression in *TP53*-mutant THP-1 cells (**Fig. 3C, top panel**). Similarly, MCL-1 and BCL-2 protein levels were downregulated in response to revumenib in *TP53* WT MV-4-11 cells, but not in *TP53* KO MV-4-11 cells (**Fig. 3C, bottom panel**), whereas other BH3 proteins were unchanged (**Fig. S4**). Based on these data, we asked if the MCL-1 inhibitor MIK665 or the BCL-2 inhibitor venetoclax might restore the apoptotic response of *TP53*-mutant AML cells to revumenib. Remarkably, we observed potent synergy when revumenib was combined with MIK665 in *TP53* KO MV-4-11 cells that surpassed the response to combination treatment in *TP53* WT MV-4-11 cells (**Figs. 3D and 3E**). Conversely, while venetoclax and revumenib synergistically induced apoptosis in *TP53* WT MV-4-11 cells, this combination was less effective in *TP53* KO MV-4-11 cells (**Figs. 3F and 3G**).

## DISCUSSION

While *RAS* mutations are the most common secondary alteration in *KMT2A*-rearranged acute leukemias, recent studies suggest that *TP53* mutations also occur frequently^10,19^. Furthermore, as the clinical development of menin inhibitors accelerates as monotherapy and in combination with chemotherapy and other targeted drugs, identifying patients who are more likely to respond and what combinations are more likely to be effective in different genetic contexts are key questions.

Fiskus, et al. reported synergistic growth inhibition of AML cell lines treated with revumenib in combination with venetoclax and the FLT3 inhibitor gilteritinib^19^. These investigators also inactivated *TP53* in MOLM-13 cells, but could not reliably ascertain how this affected the response to revumenib because unmodified, *TP53* WT MOLM-13 cells did not undergo apoptosis. We confirmed that MOLM-13 cells display intrinsic resistance to revumenib (**Fig. S5**) and went on to generate isogenic WT and *TP53*-mutant MV-4-11 cells to address the effects of *TP53* inactivation on revumenib sensitivity in AML cells that are responsive to menin inhibition at baseline. We further show that the synergistic anti-leukemia activity of revumenib and venetoclax is blunted in *TP53* KO AML cells.

By contrast, *TP53* KO AML cells are unexpectedly more sensitive than WT cells to combination treatment with revumenib and the MCL-1 inhibitor MIK665. These results align with previous studies that demonstrated the combination of MCL-1 inhibitors with venetoclax potentiated each another in leukemia^20,21^ and lung cancer^22^. However, the combination of BCL-2 and MCL-1 inhibitors is unlikely to be tolerated due to overlapping toxicity profiles, whereas strategies targeting orthogonal pathways, such as combining MCL-1 and menin inhibitors may provide a more tolerable and effective therapeutic strategy in *KMT2A*-rearranged leukemias.

In summary, we provide the first evidence that *TP53* mutations confer *de novo* resistance to menin inhibitors in *KMT2A*-rearranged AML. Accordingly, it will be important to analyze data from ongoing clinical trials to determine if *TP53* mutational status is a biomarker of intrinsic treatment resistance in *KMT2A*-rearranged AML. Furthermore, our studies provide a rationale for evaluating MCL-1 and menin inhibitor combinations in patients with treatment-refractory *TP53*-mutant and *KMT2A*-rearranged leukemias.

## METHODS

### Cell line authentication and quality control

Cell lines were obtained from ATCC or DSMZ. Cells were cultured in RPMI media containing 10% heat inactivated FBS, 1% penicillin-streptomycin and 1% GlutaMAX™ at 37ºC and in 5% CO2. Cell lines were regularly tested for mycoplasma and their identity confirmed via STR profiling.

### Cell viability and apoptosis analysis

Cells were plated at 5,000 cells/well in 96-well plates and were treated for 72 hours. Metabolically active cells as a proxy for overall cell number were measured with CellTiter-Glo (Promega). To assess for apoptosis, cells were plated at 25,000 cells/well in 96-well plates and treated for 48 hours. Cells were then fixed with 4% paraformaldehyde for 10 minutes at RT, permeabilized with methanol for 30 minutes at 4ºC, and stained with cleaved caspase-3 antibody (BD 560627) for one hour before analyzing on a flow cytometer. Experiments were performed in technical triplicate and a representative experiment is shown from at least three experimental replicates.

### Immunoblotting

Immunoblotting was performed using antibodies against BCL-2 (CST 3498S), MCL-1 (CST 5453), and HSP-90 (BD 610418) as a loading control.

## ACKNOWLEDGEMENTS

This work was supported by grants from the National Institutes of Health (K08 CA256489 [B.J.H.], R01 CA193994 [K.S.], and U54 CA196519 [K.S.]). We are grateful to the members of the Helen Diller Family Comprehensive Cancer Center core facilities in Computational Biology and Informatics and Laboratory for Cell Analysis (P30 CA082103) and the Gladstone Institutes Genomics Core. The results published here are in part based on data generated by the Therapeutically Applicable Research to Generate Effective Treatments (TARGET) project managed by the National Cancer Institute. Information about TARGET can be found at https://ocg.cancer.gov.

## AUTHORSHIP CONTRIBUTION

Conceptualization, J.D., B.J.H.; Methodology, J.D., C.J.L., B.S.B., B.J.H.; Investigation, J.D., C.J.L., A.C.B., A.S.L., A.B.; Formal Analysis, J.D., C.J.L., P.G., L.F., X.M.; Resources, A.J.L., R.E.R., C.E.M., C.C., B.S.B.; Writing, J.D., B.J.H.; Revisions, all authors; Supervision, X.M., K.S., S.M., B.S.B., B.J.H.; Funding Acquisition, K.S., S.M., B.J.H.

## DATA AVAILABILITY STATEMENT

Raw sequencing FASTQ files are available at NCBI Gene Expression Omnibus (GEO) and the NCBI Database of Genotypes and Phenotypes (dbGaP: https://www.ncbi.nlm.nih.gov/gap under study ID phs000465.v21.p8 for human bulk RNA sequencing data).

