## Supplemental Figures for "*TP53* Inactivation Confers Resistance to the Menin Inhibitor Revumenib in Acute Myeloid Leukemia"

### Supplemental Figure 1

| Table S3. Fusion Calls From RNA sequencing | | | | |
|---|---|---|---|---|
| Cell Line | Fusion or Mutation | Sequence ID | Exons | Breakpoint Sequence |
| MV-4-11 | KMT2A::AFF1 | NM_001197104 :: NM_001166693 | E8::E6 | ATCCCTGTAAAACAAAAACCAAAAGAAAAG :: GAAATGACCCATTCATGGCCGCCTCCTTTG |
| THP-1 | KMT2A::MLLT3 | NM_001197104 :: NM_004529 | E8::E6 | ATCCCTGTAAAACAAAAACCAAAAGAAAAG :: TCTGAACAACCCAGTCCTGCCAGCTCCAGC |
| NOMO-1 | KMT2A::MLLT3 | NM_001197104 :: NM_004529 | E9::E6 | GGAGTCCACAGGATCAGAGTGGACTTTAAG :: TCTGAACAACCCAGTCCTGCCAGCTCCAGC |

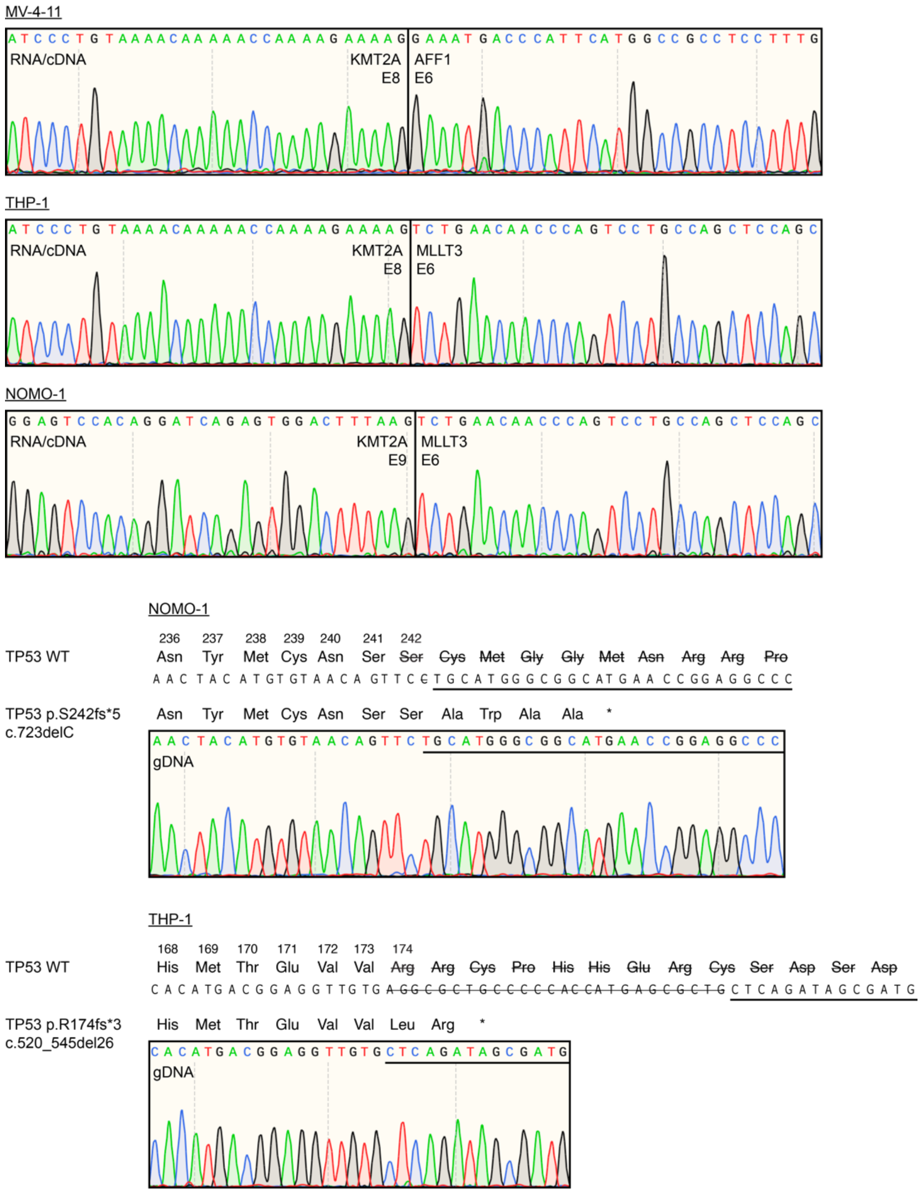


**Figure S1.** Cell line fusion and *TP53* mutation validation.

**Supplemental Figure 2**

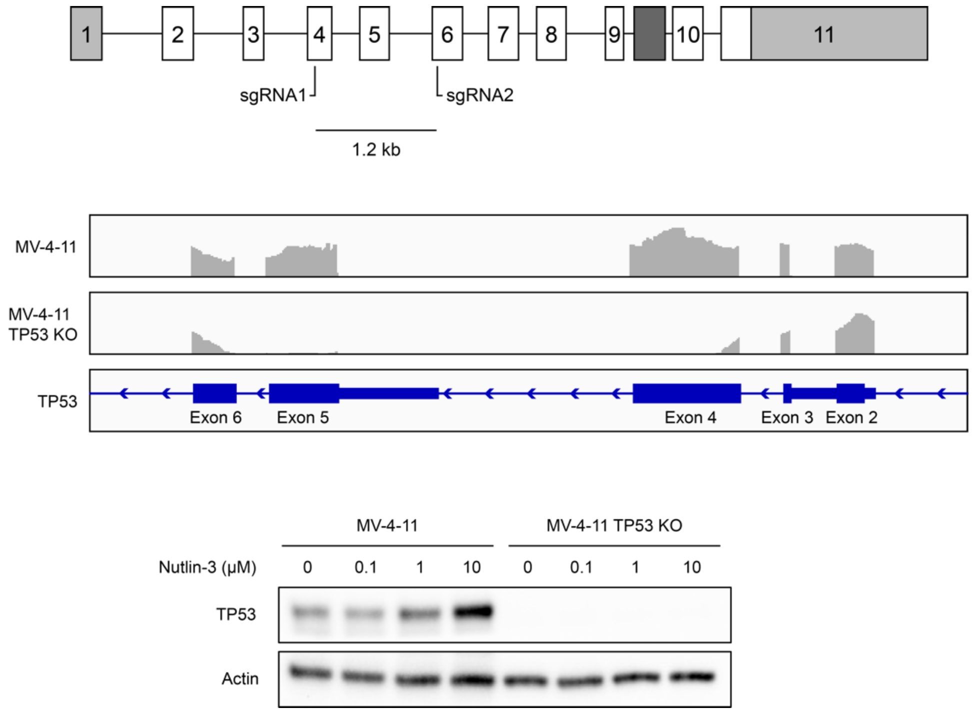


**Figure S2.** *TP53* knockout strategy. Two sgRNAs that target *TP53* for CRISPR/Cas9 editing were used together to generate a 1.2 kb deletion between the two sgRNA sites (top panel). Single cell clones were isolated by limit dilution and screened by PCR. Homozygous clones were validated by RNA-seq (middle panel) and Western blot (bottom panel).

**Supplemental Figure 3**

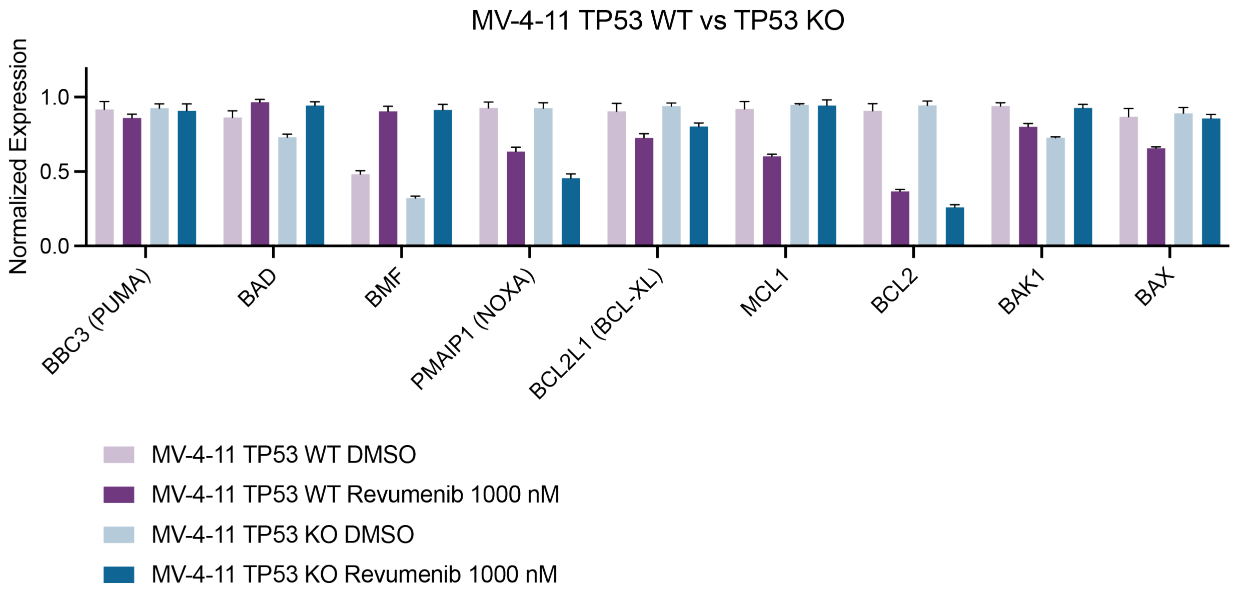


**Figure S3.** Normalized gene expression of BH3 family members in MV-4-11 TP53 wildtype (WT) versus MV-4-11 TP53 knockout (KO) cells treated with 0 and 1000 nM of revumenib for 24 hours.

**Supplemental Figure 4**

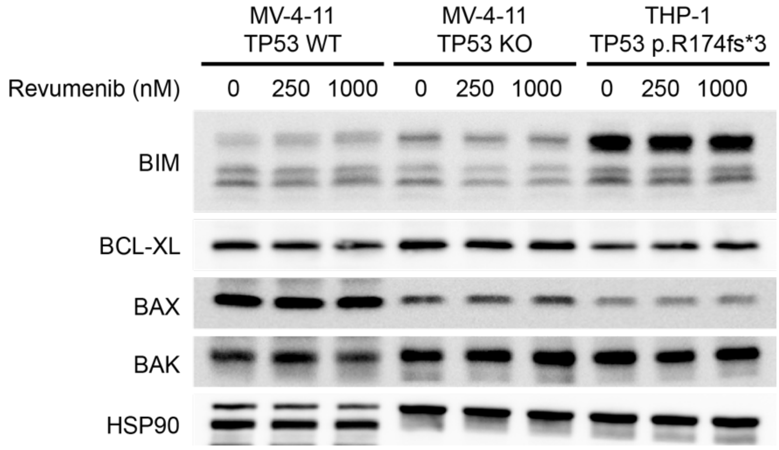


**Figure S4.** Western blot of BH3 proteins in MV-4-11 TP53 wildtype (WT), MV-4-11 TP53 knockout (KO), and THP-1 cells treated with 0, 250, and 1000 nM of revumenib for 48 hours.

**Supplemental Figure 5**

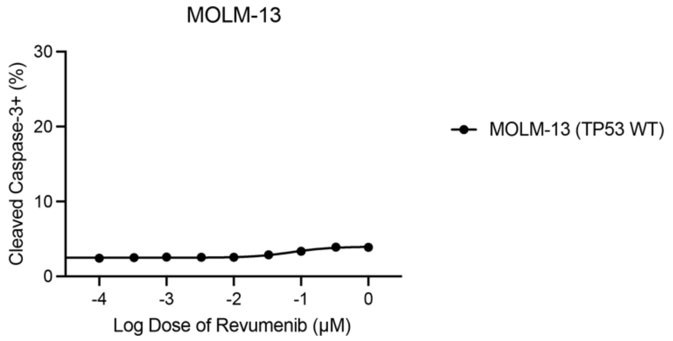


**Figure S5.** Percentage of cells positive for cleaved caspase-3 with MOLM-13 (*TP53* wildtype) cells after exposure to single agent revumenib.
